# *But Mouse, you are not alone*: On some severe acute respiratory syndrome coronavirus 2 variants infecting mice

**DOI:** 10.1101/2021.08.04.455042

**Authors:** Michael J. Kuiper, Laurence OW Wilson, Shruthi Mangalaganesh, Carol Lee, Daniel Reti, Seshadri S Vasan

**Affiliations:** Biomolecular Modeller who leads the Modelling and Simulations Team in at the CSIRO Data61, Dockands, Victoria, Australia; Research Scientist who leads the Digital Genome Engineering Team at the CSIRO Transformational Bioinformatics Group, North Ryde, New South Wales, Australia; Bachelor of Biomedical Science and Bachelor of Commerce at Monash University, Clayton, Victoria, Australia, and pursued an internship at the CSIRO Australian Centre for Disease Preparedness, Geelong, Victoria, Australia; CERC Postdoctoral Research Fellow in the CSIRO Transformational Bioinformatics Group, North Ryde, New South Wales, Australia; Research Technician at the CSIRO Transformational Bioinformatics Group, North Ryde, New South Wales, Australia, and is currently Front-End Software Engineer at the Centre for Population Genomics, joint initiative of the Garvan Institute of Medical Research, Sydney, New South Wales, Australia and the Murdoch Children’s Research Institute, Melbourne, Victoria, Australia; Senior Principal Research Consultant and COVID-19 Project Leader at the CSIRO Australian Centre for Disease Preparedness, Geelong, Victoria, Australia, and Honorary Professor in the Department of Health Sciences at the University of York, York, UK

**Keywords:** *AlphaFold*, Animal reservoir, COVID-19, *in silico*, *in vitro*, *in vivo*, mouse adaptation, SARS-CoV-2, variants

## Abstract

*In silico* predictions combined with *in vitro, in vivo* and *in situ* observations collectively suggest that mouse adaptation of the SARS-CoV-2 virus requires an aromatic substitution in position 501 or position 498 (but not both) of the spike protein’s receptor binding domain. This effect could be enhanced by mutations in positions 417, 484, and 493 (especially K417N, E484K, Q493K and Q493R), and to a lesser extent by mutations in positions 486 and 499 (such as F486L and P499T). Such enhancements due to more favourable binding interactions with residues on the complementary angiotensin-converting enzyme 2 (ACE2) interface, are however, unlikely to sustain mouse infectivity on their own based on theoretical and experimental evidence to date. Our current understanding thus points to the Alpha, Beta, Gamma, and Omicron variants of concern infecting mice, while Delta and ‘Delta Plus’ lack a similar biomolecular basis to do so. This paper identifies eleven countries (Brazil, Chile, Djibouti, Haiti, Malawi, Mozambique, Reunion, Suriname, Trinidad and Tobago, Uruguay and Venezuela) where targeted local field surveillance of mice is encouraged because they may have come in contact with humans who had the virus with adaptive mutation(s). It also provides a systematic methodology to analyze the potential for other animal reservoirs and their likely locations.

## Introduction

The ‘novel coronavirus disease’ (COVID-19) has resulted in significant global morbidity and mortality on a scale similar to the influenza pandemic of 1918.^1^ The ongoing pandemic has been sustained through the activities of human beings, who are the largest reservoir of the causative ‘severe acute respiratory syndrome virus 2’ (SARS-CoV-2). This RNA virus in a new host (human beings) is evolving rapidly, accumulating mutations, and existing as a cloud of variants with quasispecies diversity.^2^ Last year, the world witnessed the risk of this virus acquiring additional reservoirs (such as minks) and new mutations of consequence (such as ‘Cluster 5’), which could increase transmissibility and lead to a potentially weaker antibody response.^3^

Both the original form of the virus (known as ‘D614’) and the subsequent more transmissible ‘G614’ variant which has replaced it almost entirely in circulation,^4,5^ did not infect mice because their ‘angiotensin-converting enzyme 2’ (ACE2) receptor did not bind the viral spike protein effectively to allow entry into cells. Since mouse (*Mus musculus*) is a popular animal model of infection, the virus had to be adapted through techniques such as sequential passaging in mouse lung tissues and modifying the receptor binding domain (RBD).^6-8^ Other strategies to infecting mice with the original form of the virus or the G614 included transgenic mice expressing human ACE2 (hACE2), and sensitizing the mouse respiratory tract through transduction with adenovirus or adeno-associated virus expressing hACE2.^9-12^ Recently, virus ‘variants of concern’ (VOC) originating from Brazil, South Africa and the UK (which contain the common mutation N501Y in the RBD) were shown to infect mice.^13,14^ This ability of SARS-CoV-2 variants of concern to infect mice is unsettling because of the potential to establish additional reservoirs in a species that is in close contact with people and companion animals, especially as its population is hard to vaccinate or control.

## Methodology

In this paper, we have combined our structural predictions from biomolecular modelling with available experimental evidence to understand how specific mutations in VOC and mouse adapted strains, especially in the RBD, have enabled this virus to infect mice. To do this, we compared the spatial interactions of these key mutations with the corresponding regions in both mouse ACE2 (mACE2) and hACE2 using homology molecular models based on cryogenic electron microscopy (cryo-EM) data of the ACE2/RBD interface (protein database ‘pdb’ entry number 6M17).^15^ We have assumed the coordinates of ACE2 and RBD in the 6M17 structure to be reasonable starting positions for our respective models, with root-mean-square deviation (RMSD) scripts alignment based on C-Alpha carbon protein backbones. As the protein prediction software *AlphaFold* has been recently been released,^16^ we also recalculated RBD structures, finding *AlphaFold* predictions to be in excellent agreement with our initial homology models, with less than 0.9 Å RMSD average alignment. Amino acid side chain conformations of the RBD models were adapted from *AlphaFold* predictions as they were deemed more reliable.

By introducing the ACE2 models and variants into molecular dynamics simulations to optimize the interactions, similar to our recent work,^17,18^ we qualitatively identified the key interactions of side chain residues at the ACE2/RBD interface. Further details are provided under ‘Supplementary Methods for Molecular Modelling’. We then compared our *in-silico* findings with experimental results reported by different research groups with mouse adapted strains/isolates^6-8,13,21-29^ and VOC^14,30-34^ (Tables 1 and 2). Taken together, we were able to gain valuable insights into the likely effects of different mutations of consequence.

**Table 1.** Key mutations in 14 ‘mouse adapted strains’ and 3 ‘variants of concern’ of SARS-CoV-2 known to infect mice, with further characterization as outlined in **Table 2**.

| Mice-infecting SARS-CoV-2 'strains'/isolates | Key mutations in the spike RBD (residues 331-524)<sup>19</sup> |  |  |  |  |  |  | Color legend – mutations scored on Grantham Distance,<sup>20</sup> in the scale of 5 (conservative) to 215 (radical), similar to Riddell et al.<sup>18</sup> |  |  |  |  |  |  |  |  |  |  |
| --- | --- | --- | --- | --- | --- | --- | --- | --- | --- | --- | --- | --- | --- | --- | --- | --- | --- | --- |
|  | K417 | E484 | F486 | Q493 | Q498 | P499 | N501 | 22 | 24 | 38 | 43 | 53 | 56 | 73 | 94 | 95 | 99 | 143 |
| Mouse Adapted (MA) strains |  |  |  |  |  |  |  | Authors' synthesis/comments |  |  |  |  |  |  |  |  |  |  |
| MASCP6<sup>6</sup> |  |  |  |  |  |  | N→Y | L37F (nsp6), P84S (nsp10), and D128Y (N) were the other observed nonsynonymous mutations. |  |  |  |  |  |  |  |  |  |  |
| MASCP25<sup>21</sup> |  |  |  | Q→K |  |  | N→Y | MASCP6 passaged serially to develop MASCP25 and MASCP36, which accumulated Q493K and subsequently K417N in the RBD. |  |  |  |  |  |  |  |  |  |  |
| MASCP36<sup>21</sup> | K→N |  |  | Q→K |  |  | N→Y | Highly virulent in aged mice and reported stable/mature RBD binding. It also had I1258V (nsp3), S301L (nsp5), and R32C (N). |  |  |  |  |  |  |  |  |  |  |
| IC-MA1<sup>7</sup> |  |  |  |  | Q→Y | P→T |  | Spike Q498Y and P499T were engineered for compatibility with Q42 of mouse ACE2. |  |  |  |  |  |  |  |  |  |  |
| IC-MA10<sup>8</sup> |  |  |  | Q→K | Q→Y | P→T |  | Q493K evolved in MA10 and interacts with mACE2's N31; others were T285I (nsp4), K2R (nsp7), E23G (nsp8), and F7S (ORF6). |  |  |  |  |  |  |  |  |  |  |
| WA1-MA-P11<sup>13</sup> |  |  |  |  |  |  | N→Y | In addition to N501Y, there were H655Y, T7I (M), L84S (ORF8), S194T (N) and the +KLRS (216-219) insertion also observed by others and predicted to form a solvent-accessible loop.<sup>22</sup> |  |  |  |  |  |  |  |  |  |  |
| HRB26M<sup>23</sup> (HRB26-P14) |  |  |  |  | Q→H |  |  | HRB26-P5 had Q498H, N969S, and the 675QTQTN679 deletion observed by others.<sup>24,25</sup> P14 also had A81T (nsp8). |  |  |  |  |  |  |  |  |  |  |
| Hu-1-WBP-1<sup>26</sup> |  |  |  | Q→K | Q→H |  |  | Q493K and Q498H by P5, interacting respectively with mACE2's N31 via H-bond and Y41 via π-stacking. Nonsynonymous Q493K and T77I (nsp9) mutations increased frequency from P8 to P11. |  |  |  |  |  |  |  |  |  |  |
| IC-MA4<sup>27</sup> |  |  | F→L |  | Q→Y |  |  | F486L and Q498Y were engineered for infecting mice. |  |  |  |  |  |  |  |  |  |  |
| IC-CMA1-3<sup>27</sup> | K→N |  |  |  |  |  | N→Y | CMA1-3 all had N501Y. CMA2 also had D128Y; CMA3 also had D128Y, L37F, and P87S. Passaging of CMA3 resulted in K417N, H655Y, and E8V (E). |  |  |  |  |  |  |  |  |  |  |
| LG<sup>28</sup> |  |  |  |  | Q→H |  |  | Other mutations present were N74K, H655Y, I6V (ORF10). |  |  |  |  |  |  |  |  |  |  |
| IA-N501Y-MA30 | K→M | E→K |  | Q→R | Q→R |  | N→Y | Homogenate from BALB/c mouse lung after 30 passages. GISAID entry EPI_ISL_1666328.<sup>29</sup> |  |  |  |  |  |  |  |  |  |  |
| Variants of Concern (VOC) infecting mice |  |  |  |  |  |  |  |  |  |  |  |  |  |  |  |  |  |  |
| Alpha (501Y.V1 or B.1.1.7<sup>14,30</sup>) |  | E→K |  |  |  |  | N→Y | First reported in the UK, September 2020. Spike also has A570D, P681H, T716I, S982A, and D1118H, in addition to HV69-70 and Y144 deletions. **E484K has been detected in this VOC in 15 countries as of 29 April 2021 (436 entries on GISAID).** |  |  |  |  |  |  |  |  |  |  |
| Beta (501Y.V2 or B.1.351<sup>14,31,32</sup>) | K→N | E→K |  |  |  |  | N→Y | First reported in South Africa, May 2020. V2-1 had D80A, D215G, E484K, N501Y, D614G and A701V in the Spike. L18F and K417N arose in V2-2. V2-3 had 242-244 deletion. |  |  |  |  |  |  |  |  |  |  |
| Gamma (501Y.V3 or P.1<sup>14,32,33</sup>) | K→T | E→K |  |  |  |  | N→Y | First reported in Brazil, November 2020. L18F, T20N, P26S, D138Y, R190S, K417T, E484K, N501Y, H655Y and T1027I in Spike. |  |  |  |  |  |  |  |  |  |  |
| VOC not known to infect mice |  |  |  |  |  |  |  |  |  |  |  |  |  |  |  |  |  |  |
| Delta (452R.V3 or B.1.617.2)<sup>34</sup> | K→N |  |  |  |  |  |  | First reported in India, October 2020. Also has L452R in the RBD; G142D, R158G, T478K, P681R and D950N in Spike, along with deletions 156-157. **K417N has been detected in this VOC (called 'Delta Plus') in 12 countries as of 25 June 2021.** The related Kappa variant of interest (B.1.617.1) has L452R and E484Q in RBD (Grantham Distance scores of 102 and 29 respectively). |  |  |  |  |  |  |  |  |  |  |

**Table 2.**
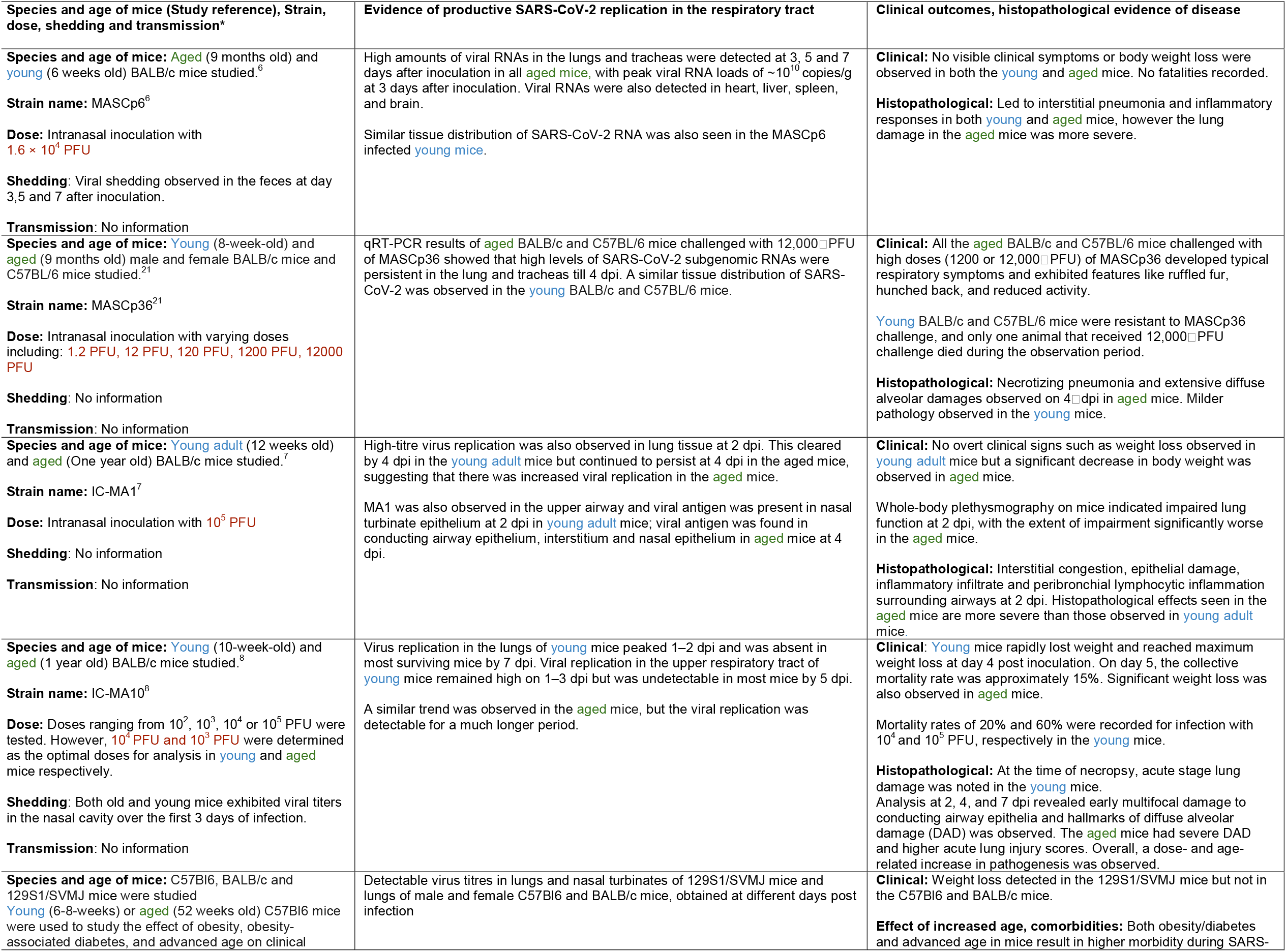
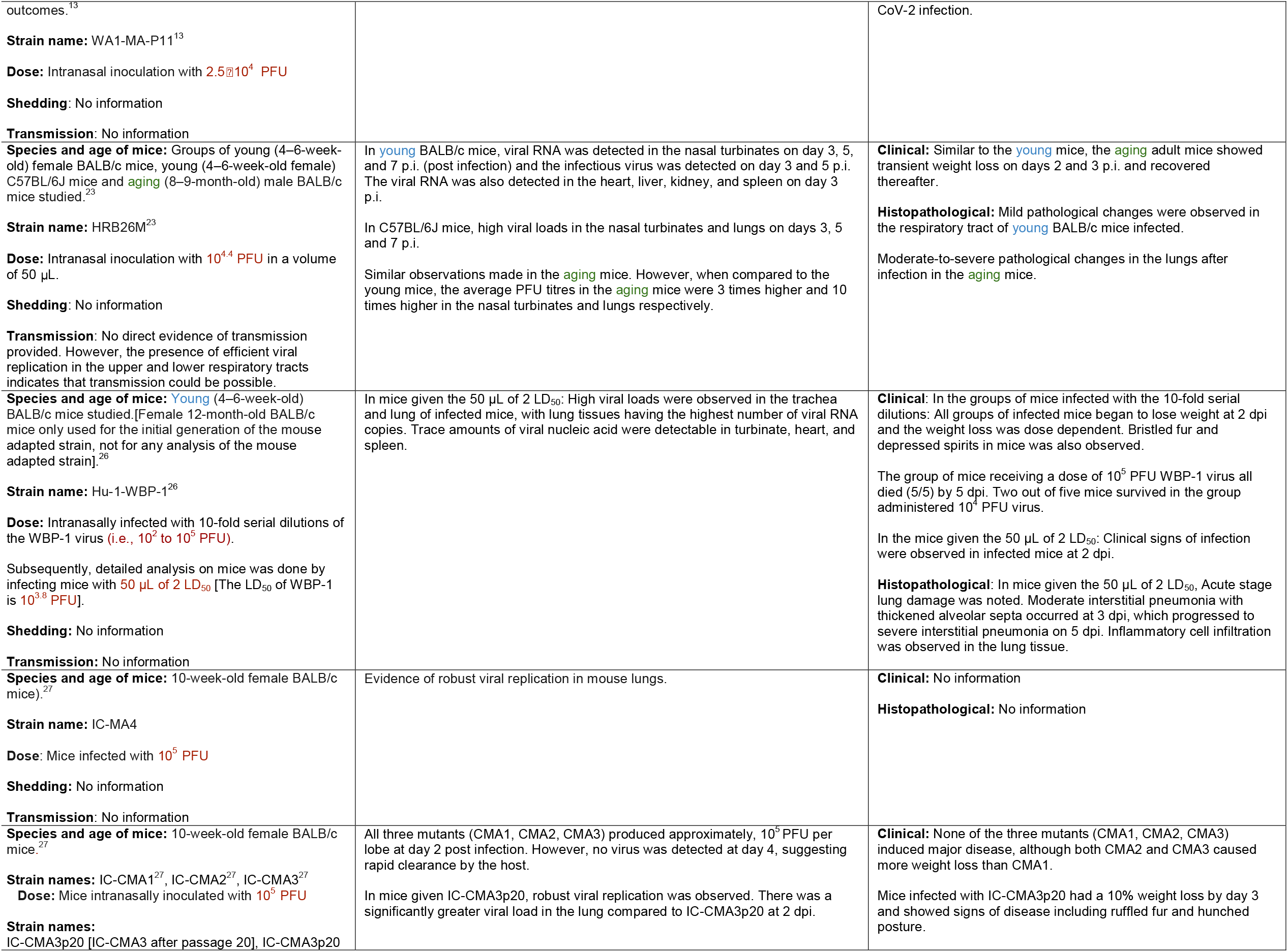
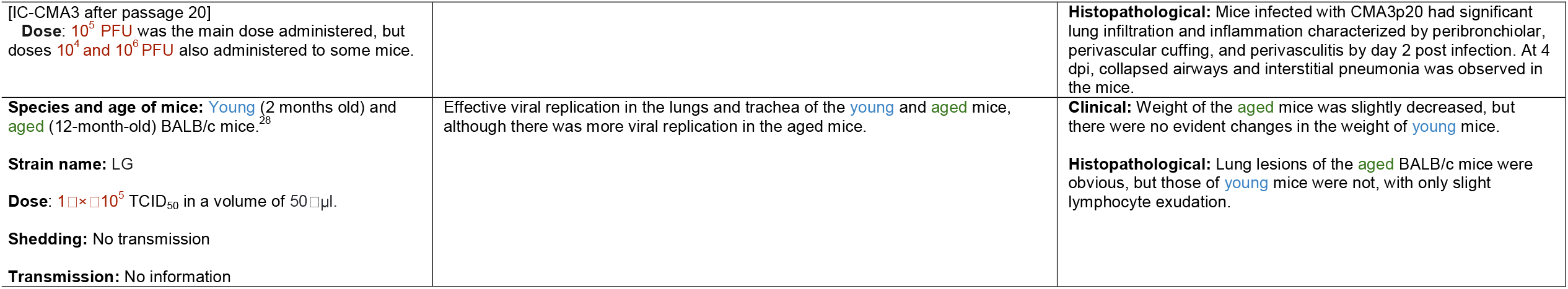
Description of the characteristics of the mouse adapted strains generated from the *in vivo* studies discussed in **Table 1**. *Strain = Mouse adapted strain name; Dose = Dose of SARS-CoV-2 mouse adapted strain provided to the mice stated in plaque forming units (PFU) or the tissue culture infective Dose (TCID50); Shedding & Transmission = Any evidence of viral shedding and viral transmission observed in the mice.

| Species and age of mice (Study reference), Strain, dose, shedding and transmission* | Evidence of productive SARS-CoV-2 replication in the respiratory tract | Clinical outcomes, histopathological evidence of disease |
| --- | --- | --- |
| **Species and age of mice:** **Aged** (9 months old) and **young** (6 weeks old) BALB/c mice studied.<sup>6</sup><br>**Strain name:** MASCp6<sup>6</sup><br>**Dose:** Intranasal inoculation with **1.6 × 10<sup>4</sup> PFU**<br>**Shedding:** Viral shedding observed in the feces at day 3,5 and 7 after inoculation.<br>**Transmission:** No information | High amounts of viral RNAs in the lungs and tracheas were detected at 3, 5 and 7 days after inoculation in all **aged mice**, with peak viral RNA loads of ~10<sup>10</sup> copies/g at 3 days after inoculation. Viral RNAs were also detected in heart, liver, spleen, and brain.<br>Similar tissue distribution of SARS-CoV-2 RNA was also seen in the MASCp6 infected **young mice**. | **Clinical:** No visible clinical symptoms or body weight loss were observed in both the **young** and **aged** mice. No fatalities recorded.<br>**Histopathological:** Led to interstitial pneumonia and inflammatory responses in both **young** and **aged** mice, however the lung damage in the **aged** mice was more severe. |
| **Species and age of mice:** **Young** (8-week-old) and **aged** (9 months old) male and female BALB/c mice and C57BL/6 mice studied.<sup>21</sup><br>**Strain name:** MASCp36<sup>21</sup><br>**Dose:** Intranasal inoculation with varying doses including: **1.2 PFU, 12 PFU, 120 PFU, 1200 PFU, 12000 PFU**<br>**Shedding:** No information<br>**Transmission:** No information | qRT-PCR results of **aged** BALB/c and C57BL/6 mice challenged with 12,000□PFU of MASCp36 showed that high levels of SARS-CoV-2 subgenomic RNAs were persistent in the lung and tracheas till 4 dpi. A similar tissue distribution of SARS-CoV-2 was observed in the **young** BALB/c and C57BL/6 mice. | **Clinical:** All the **aged** BALB/c and C57BL/6 mice challenged with high doses (1200 or 12,000□PFU) of MASCp36 developed typical respiratory symptoms and exhibited features like ruffled fur, hunched back, and reduced activity.<br>**Young** BALB/c and C57BL/6 mice were resistant to MASCp36 challenge, and only one animal that received 12,000□PFU challenge died during the observation period.<br>**Histopathological:** Necrotizing pneumonia and extensive diffuse alveolar damages observed on 4□dpi in **aged** mice. Milder pathology observed in the **young** mice. |
| **Species and age of mice:** **Young adult** (12 weeks old) and **aged** (One year old) BALB/c mice studied.<sup>7</sup><br>**Strain name:** IC-MA1<sup>7</sup><br>**Dose:** Intranasal inoculation with **10<sup>5</sup> PFU**<br>**Shedding:** No information<br>**Transmission:** No information | High-titre virus replication was also observed in lung tissue at 2 dpi. This cleared by 4 dpi in the **young adult** mice but continued to persist at 4 dpi in the aged mice, suggesting that there was increased viral replication in the **aged** mice.<br>MA1 was also observed in the upper airway and viral antigen was present in nasal turbinate epithelium at 2 dpi in **young adult** mice; viral antigen was found in conducting airway epithelium, interstitium and nasal epithelium in **aged** mice at 4 dpi. | **Clinical:** No overt clinical signs such as weight loss observed in **young adult** mice but a significant decrease in body weight was observed in **aged** mice.<br>Whole-body plethysmography on mice indicated impaired lung function at 2 dpi, with the extent of impairment significantly worse in the **aged** mice.<br>**Histopathological:** Interstitial congestion, epithelial damage, inflammatory infiltrate and peribronchial lymphocytic inflammation surrounding airways at 2 dpi. Histopathological effects seen in the **aged** mice are more severe than those observed in **young adult** mice. |
| **Species and age of mice:** **Young** (10-week-old) and **aged** (1 year old) BALB/c mice studied.<sup>8</sup><br>**Strain name:** IC-MA10<sup>8</sup><br>**Dose:** Doses ranging from 10<sup>2</sup>, 10<sup>3</sup>, 10<sup>4</sup> or 10<sup>5</sup> PFU were tested. However, **10<sup>4</sup> PFU and 10<sup>3</sup> PFU** were determined as the optimal doses for analysis in **young** and **aged** mice respectively.<br>**Shedding:** Both old and young mice exhibited viral titers in the nasal cavity over the first 3 days of infection.<br>**Transmission:** No information | Virus replication in the lungs of **young** mice peaked 1–2 dpi and was absent in most surviving mice by 7 dpi. Viral replication in the upper respiratory tract of **young** mice remained high on 1–3 dpi but was undetectable in most mice by 5 dpi.<br>A similar trend was observed in the **aged** mice, but the viral replication was detectable for a much longer period. | **Clinical:** **Young** mice rapidly lost weight and reached maximum weight loss at day 4 post inoculation. On day 5, the collective mortality rate was approximately 15%. Significant weight loss was also observed in **aged** mice.<br>Mortality rates of 20% and 60% were recorded for infection with 10<sup>4</sup> and 10<sup>5</sup> PFU, respectively in the **young** mice.<br>**Histopathological:** At the time of necropsy, acute stage lung damage was noted in the **young** mice. Analysis at 2, 4, and 7 dpi revealed early multifocal damage to conducting airway epithelia and hallmarks of diffuse alveolar damage (DAD) was observed. The **aged** mice had severe DAD and higher acute lung injury scores. Overall, a dose- and age-related increase in pathogenesis was observed. |
| **Species and age of mice:** C57Bl6, BALB/c and 129S1/SVMJ mice were studied<br>**Young** (6-8-weeks) or **aged** (52 weeks old) C57Bl6 mice were used to study the effect of obesity, obesity-associated diabetes, and advanced age on clinical | Detectable virus titres in lungs and nasal turbinates of 129S1/SVMJ mice and lungs of male and female C57Bl6 and BALB/c mice, obtained at different days post infection | **Clinical:** Weight loss detected in the 129S1/SVMJ mice but not in the C57Bl6 and BALB/c mice.<br>**Effect of increased age, comorbidities:** Both obesity/diabetes and advanced age in mice result in higher morbidity during SARS- |
| outcomes.<sup>13</sup><br>**Strain name:** WA1-MA-P11<sup>13</sup><br>**Dose:** Intranasal inoculation with **2.5×10<sup>4</sup> PFU**<br>**Shedding:** No information<br>**Transmission:** No information |  | CoV-2 infection. |
| **Species and age of mice:** Groups of young (4–6-week-old) female BALB/c mice, young (4–6-week-old female) C57BL/6J mice and **aging** (8–9-month-old) male BALB/c mice studied.<sup>23</sup><br>**Strain name:** HRB26M<sup>23</sup><br>**Dose:** Intranasal inoculation with **10<sup>4.4</sup> PFU** in a volume of 50 µL.<br>**Shedding:** No information<br>**Transmission:** No direct evidence of transmission provided. However, the presence of efficient viral replication in the upper and lower respiratory tracts indicates that transmission could be possible. | In **young** BALB/c mice, viral RNA was detected in the nasal turbinates on day 3, 5, and 7 p.i. (post infection) and the infectious virus was detected on day 3 and 5 p.i. The viral RNA was also detected in the heart, liver, kidney, and spleen on day 3 p.i.<br>In C57BL/6J mice, high viral loads in the nasal turbinates and lungs on days 3, 5 and 7 p.i.<br>Similar observations made in the **aging** mice. However, when compared to the young mice, the average PFU titres in the **aging** mice were 3 times higher and 10 times higher in the nasal turbinates and lungs respectively. | **Clinical:** Similar to the **young** mice, the **aging** adult mice showed transient weight loss on days 2 and 3 p.i. and recovered thereafter.<br>**Histopathological:** Mild pathological changes were observed in the respiratory tract of **young** BALB/c mice infected.<br>Moderate-to-severe pathological changes in the lungs after infection in the **aging** mice. |
| **Species and age of mice:** **Young** (4–6-week-old) BALB/c mice studied.[Female 12-month-old BALB/c mice only used for the initial generation of the mouse adapted strain, not for any analysis of the mouse adapted strain].<sup>26</sup><br>**Strain name:** Hu-1-WBP-1<sup>26</sup><br>**Dose:** Intranasally infected with 10-fold serial dilutions of the WBP-1 virus (**i.e., 10<sup>2</sup> to 10<sup>5</sup> PFU**).<br>Subsequently, detailed analysis on mice was done by infecting mice with **50 µL of 2 LD<sub>50</sub>** [The LD<sub>50</sub> of WBP-1 is **10<sup>3.8</sup> PFU**].<br>**Shedding:** No information<br>**Transmission:** No information | In mice given the 50 µL of 2 LD<sub>50</sub>: High viral loads were observed in the trachea and lung of infected mice, with lung tissues having the highest number of viral RNA copies. Trace amounts of viral nucleic acid were detectable in turbinate, heart, and spleen. | **Clinical:** In the groups of mice infected with the 10-fold serial dilutions: All groups of infected mice began to lose weight at 2 dpi and the weight loss was dose dependent. Bristled fur and depressed spirits in mice was also observed.<br>The group of mice receiving a dose of 10<sup>5</sup> PFU WBP-1 virus all died (5/5) by 5 dpi. Two out of five mice survived in the group administered 10<sup>4</sup> PFU virus.<br>In the mice given the 50 µL of 2 LD<sub>50</sub>: Clinical signs of infection were observed in infected mice at 2 dpi.<br>**Histopathological:** In mice given the 50 µL of 2 LD<sub>50</sub>, Acute stage lung damage was noted. Moderate interstitial pneumonia with thickened alveolar septa occurred at 3 dpi, which progressed to severe interstitial pneumonia on 5 dpi. Inflammatory cell infiltration was observed in the lung tissue. |
| **Species and age of mice:** 10-week-old female BALB/c mice).<sup>27</sup><br>**Strain name:** IC-MA4<br>**Dose:** Mice infected with **10<sup>5</sup> PFU**<br>**Shedding:** No information<br>**Transmission:** No information | Evidence of robust viral replication in mouse lungs. | **Clinical:** No information<br>**Histopathological:** No information |
| **Species and age of mice:** 10-week-old female BALB/c mice.<sup>27</sup><br>**Strain names:** IC-CMA1<sup>27</sup>, IC-CMA2<sup>27</sup>, IC-CMA3<sup>27</sup><br>**Dose:** Mice intranasally inoculated with **10<sup>5</sup> PFU**<br>**Strain names:** IC-CMA3p20 [IC-CMA3 after passage 20], IC-CMA3p20 | All three mutants (CMA1, CMA2, CMA3) produced approximately, 10<sup>5</sup> PFU per lobe at day 2 post infection. However, no virus was detected at day 4, suggesting rapid clearance by the host.<br>In mice given IC-CMA3p20, robust viral replication was observed. There was a significantly greater viral load in the lung compared to IC-CMA3p20 at 2 dpi. | **Clinical:** None of the three mutants (CMA1, CMA2, CMA3) induced major disease, although both CMA2 and CMA3 caused more weight loss than CMA1.<br>Mice infected with IC-CMA3p20 had a 10% weight loss by day 3 and showed signs of disease including ruffled fur and hunched posture. |
| [IC-CMA3 after passage 20]<br>**Dose:** 10<sup>5</sup> PFU was the main dose administered, but doses 10<sup>4</sup> and 10<sup>6</sup> PFU also administered to some mice. |  | **Histopathological:** Mice infected with CMA3p20 had significant lung infiltration and inflammation characterized by peribronchiolar, perivascular cuffing, and perivascularitis by day 2 post infection. At 4 dpi, collapsed airways and interstitial pneumonia was observed in the mice. |
| **Species and age of mice:** Young (2 months old) and aged (12-month-old) BALB/c mice.<sup>28</sup><br>**Strain name:** LG<br>**Dose:** 1 × 10<sup>5</sup> TCID<sub>50</sub> in a volume of 50 µl.<br>**Shedding:** No transmission<br>**Transmission:** No information | Effective viral replication in the lungs and trachea of the young and aged mice, although there was more viral replication in the aged mice. | **Clinical:** Weight of the aged mice was slightly decreased, but there were no evident changes in the weight of young mice.<br>**Histopathological:** Lung lesions of the aged BALB/c mice were obvious, but those of young mice were not, with only slight lymphocyte exudation. |

Subsequently, we queried the world’s largest public database called ‘GISAID’ (Global Initiative on Sharing All Influenza Data) and looked for these mutations, and equivalent mutations of consequence. This ‘Big Data’, comprising of circa 2.4 million and 4 million SARS-CoV-2 genome sequences as of 21 July and 4 October 2021 respectively, formed the basis of our *in situ* analysis.^29^ Over 99.9% of these sequences are from viruses sequenced from human hosts. In brief, sequences were aligned to the back to the SARS-CoV-2 reference (EPI_ISL_402124, denoted as the WT allele) in order to generate a file containing all mutations in the variant call format (VCF), which is a concise way for storing gene sequence variations. We then calculated the ratio of the frequency at which the WT allele versus the mutant allele were observed across a rolling 14-day window. Given the data is discrete, highly variable in size across countries, and contains background noise, such a window of time is essential from our experience to reduce distortions and glean meaningful insights. For example, if 14 WT and 28 mutant sequences were observed within a specified 14-day window, then the frequencies were 1/day and 2/day respectively. The ratio of mutant:WT frequency is therefore 2, indicating the mutant allele is appearing twice as frequently as the WT allele within that period. This was used to create a heatmap for the key mutations, both individually and in combination (as discussed below). For clarity, we only considered countries where the mutant:WT ratio exceeded 1.2 across the entire time the mutant was sampled or within any given 14-day window. To reduce the noise further from low-sampled countries, we also instituted a minimum threshold of 10 WT and 10 mutant samples over at least 14 days to suggest possible spread locally. For this reason, a country where a mutant had only been recorded on a single day, or where only 9 mutant samples were recorded overall, was not included in our analysis. Our heatmap scale has also been truncated at 2.00 for ease of visual comparison.

## Results and Discussion

### *In silico* results

*In silico* comparison of the interface residues of the RBD/ACE2 complex in human and mouse models reveals 30 ACE2 residues in close contact, of which 19 are conserved between mACE2 and hACE2 (approximately 63% identity for the contacting ACE2 residues). The RBD mutations associated with the mouse adaptations listed in **Table 1** can be grouped loosely into 3 regions by their positions on the ACE2/RBD interface as follows: Region 1 (RBD positions 498, 499 and 501) are centered around the highly conserved ACE2 residue tyrosine 41 (Y41); Region 2 (RBD positions 417, 493) centered around ACE2 residue 34; and Region 3 (RBD positions 484 and 486) close to a cluster of ACE2 residues 78 to 82. ACE2 residues within 5 Å (chosen to account for molecular fluctuations) of each RBD adaptation are listed in **Table 3**, with dissimilar human/mouse ACE2 residues highlighted in yellow. **Figure 1** further illustrates these residues visually, and how they are relatively positioned in three regions. **Figure 2** aligns the human and mouse ACE2 to highlight the key differences at the contact points with the RDB (shown in yellow).

**Table 3.**
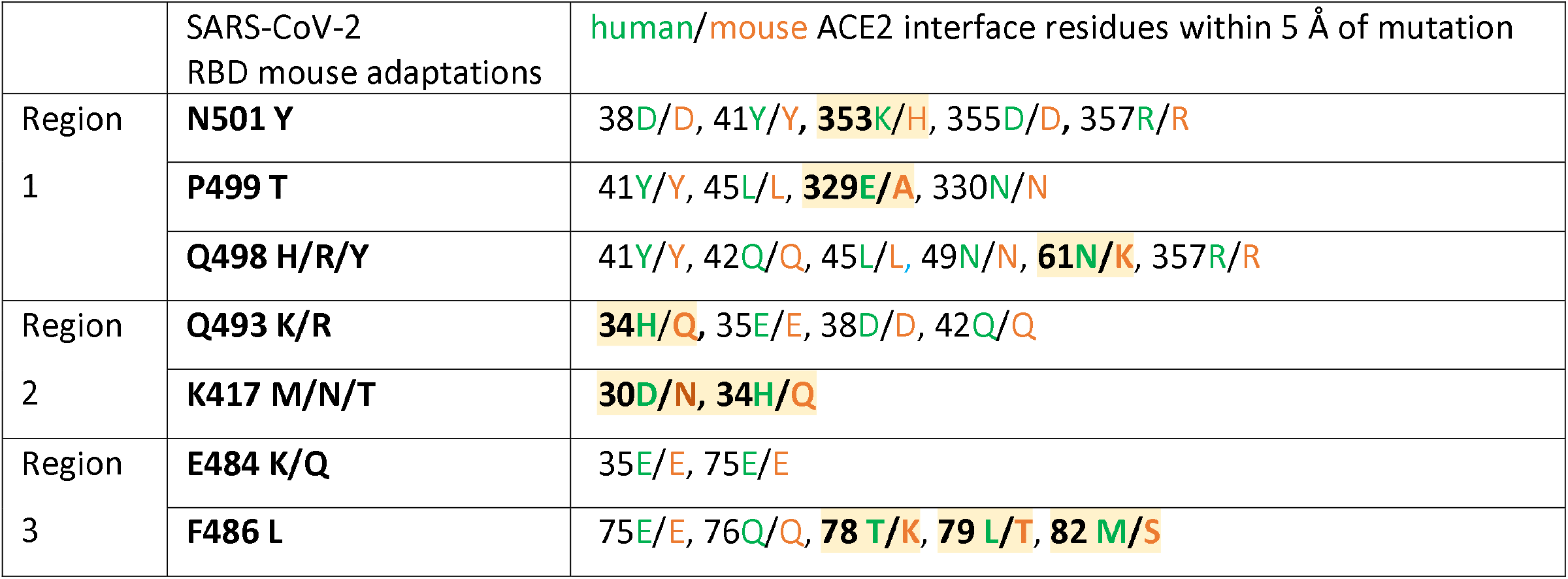
SARS-CoV-2 spike RBD mutations associated with mouse adaptation and their close contact residues in respective ACE2 proteins. Positions with dissimilar human/mouse residues are highlighted in yellow.

|  | SARS-CoV-2<br>RBD mouse adaptations | human/mouse ACE2 interface residues within 5 Å of mutation |
| --- | --- | --- |
| Region<br>1 | **N501 Y** | 38D/D, 41Y/Y, **353K/H**, 355D/D, 357R/R |
|  | **P499 T** | 41Y/Y, 45L/L, **329E/A**, 330N/N |
|  | **Q498 H/R/Y** | 41Y/Y, 42Q/Q, 45L/L, 49N/N, **61N/K**, 357R/R |
| Region<br>2 | **Q493 K/R** | **34H/Q**, 35E/E, 38D/D, 42Q/Q |
|  | **K417 M/N/T** | **30D/N**, **34H/Q** |
| Region<br>3 | **E484 K/Q** | 35E/E, 75E/E |
|  | **F486 L** | 75E/E, 76Q/Q, **78 T/K**, **79 L/T**, **82 M/S** |

**Figure 1.**
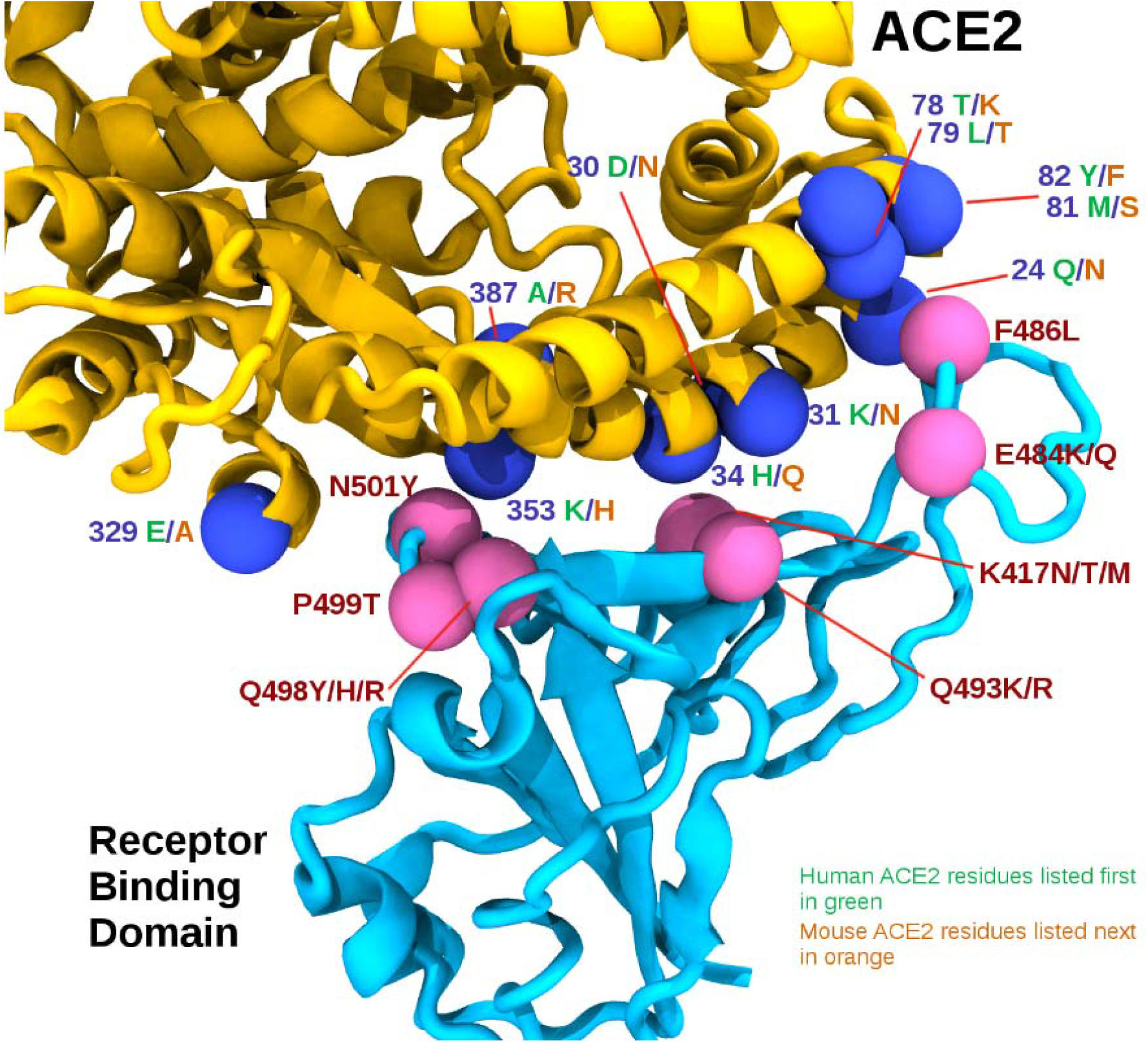
The RBD/ACE2 interface. The receptor binding domain (RBD) is shown in cyan, while the ACE2 is shown in yellow. Pink spheres indicate relative positions of mouse adapting mutations on the RBD, while the blue spheres represent interface residues that differ between human and mouse ACE2 sequences, shown in green and orange respectively.

**Figure 2.**
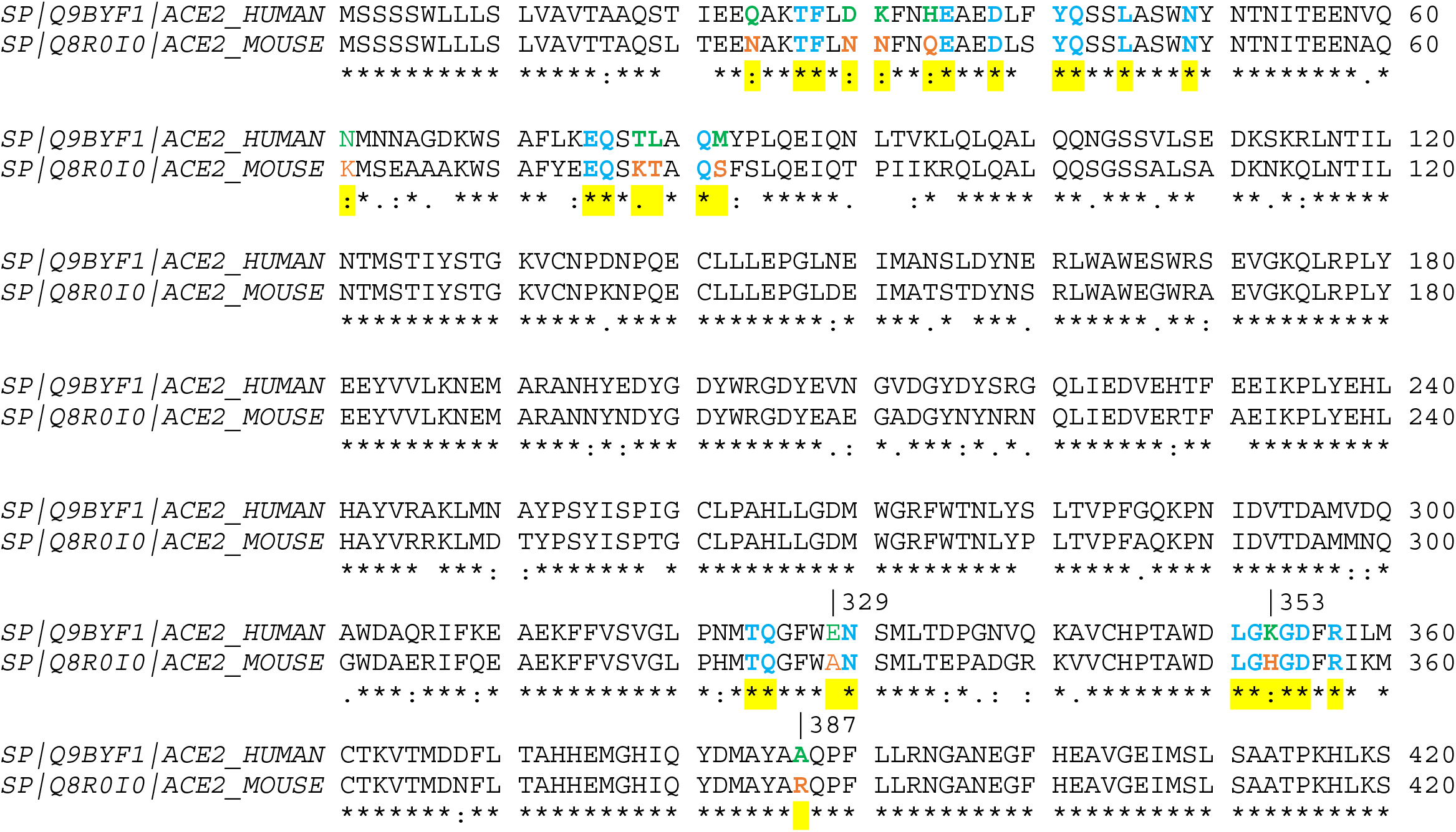
Sequence alignment of human and mouse ACE2 highlighting contact points with the SARS-CoV-2 spike receptor binding domain in yellow. Differences between contact points are highlighted in green (human) and orange (mouse), while common contact residues are highlighted in cyan.

Modelling the N501Y mutation at the ACE2/RBD interface reveals a close interaction with the highly conserved Y41 residue in ACE2, through attractive, non-covalent bonding between aromatic amino acids known as π-stacking interactions. In mACE2, π-stacking can be enhanced by the proximal substitution of histidine (H) in place of lysine (K) which is present at position 353 of hACE2. Our modelling also shows similar π-stacking enhancement with the conserved Y41, through the substitution of RBD glutamine at position 498 (Q498) to either histidine (H) or tyrosine (Y). These aromatic π-stacking interactions appear to be reasonably strong as N501Y and Q498H can each sustain mouse adaptation on its own in experiments.^6,13,23,28^ Counterintuitively, our modelling predicts that simultaneous aromatic mutations at RBD positions 498 and 501 is detrimental to mouse adaptation due to local π-stacking distortion to the binding interface. This could explain why very few simultaneous aromatic mutations at these positions have been observed. In over 2.4 million entries on GISAID as of 21 July 2021, we detected only a single co-occurrence, that of an adapted isolate from mouse lung homogenate where N501Y occurred with glutamate 498 to arginine (Q498R).^29^ However, when we recently re-run our *in situ* model with over 4 million human-origin GISAID entries, we detected 0, 2 and 14 instances of Y501 co-existing with Y498, H498 and R498 respectively. The two instances of Y501-H498 are low quality and/or low coverage sequences; the 14 instances of Y501-R498 have been reported from France, Netherlands, South Africa, Spain, UK and USA between March and September 2021. Although arginine is not aromatic, it is frequently associated with π-stacking interactions with inherent conformational flexibility compared to tyrosine or histidine, therefore it is likely that the Y501-R498 combination is more tolerated (for example, with the recent Omicron variant of concern).^37^

In Region 1 (**Table 3**) we also note the proline 499 to threonine (P499T) substitution in two infectious clones, presumably engineered to enhance Y498. Modelling provides the following insight on this: the change from P to T will relax the backbone constraints of proline and allow conformational rearrangement of threonine to contact the conserved ACE2 residues Y41 and L45. However, we don’t find any strong molecular modelling basis for this mutation to sustain mouse adaptation on its own or evolve naturally alongside adaptive mutations at positions 498 or 501; this is borne out by experimental evidence to date. In other words, *in silico* predictions combined with *in vitro* and *in vivo* evidence collectively suggest that mouse adaptation requires an aromatic substitution in either position 501 or position 498 (but not both); while additional mutations, especially in Region 2 and Region 3 of the RBD as summarized below, enhancing ACE2 binding interactions and specificity in mice. These predictions are further supported by *AlphaFold* that assigns very high confidence scores (>93 in a scale of 0-100) for the structural predictions involving these key mutations of the individual proteins, however predictions of the RBD/ACE2 complex are currently sub-optimal. With time, the quality of *AlphaFold* predictions of protein complexes will improve thanks to additional experimental observations, and thus it is expected to play a more central role in structural interpretation during pandemics such as COVID-19 and future ‘Disease-X’.

From **Table 1**, we see that mouse adapted strains sometimes carry the Q493K/R mutation (polar glutamine to basic lysine or arginine). Modelling predicts this as being enabled through favourable salt-bridge interactions with both glutamic acid 35 (E35) or aspartic acid 38 (D38) both of which are conserved in hACE2 as well as mACE2 (c.f. **Table 3**, Region 2). The K417N substitution (lysine to asparagine), which is another experimental observation from **Table 1**, is also predicted by modelling to be advantageous for mouse adaptation due to favourable amide hydrogen bond interactions with interfacial mACE2 residues asparagine 30 (N30) and glutamine 34 (Q34); such amide hydrogen bonding is not possible in hACE2 as it has non-amide lysine (K) and histidine (H) residues, as also noted by other researchers.^21^ With the Gamma variant of concern, it is unclear whether the K417T enhances the role of N501Y in mouse adaptation in a similar manner. The ‘Delta Plus’ variant of concern has the K417N mutation, but there is no molecular modelling basis to believe that it can infect mice without an aromatic change in position 498 or 501 as described above. It would be worthwhile to further investigate any interfering role of glycosylation at this interface region, because hACE2 contains N-linked glycosylation at asparagine 90 (N90) whereas mACE2 does not (its analogous residue is threonine T90 according to Uniprot references Q8R0I0 and Q9BYF1).

In Region 3, the K484 residue is not positioned directly at the interface and not observed to interact strongly with any ACE2 residues; however, our model shows occasional salt bridges can be formed with relatively close glutamic acid residues in positions 35 and 75 that are conserved in both hACE2 and mACE2. Our simulations show that the distance between K484 and these glutamic acid residues fluctuate dynamically from 3 to 20 Å, with salt bridges more likely when distances are around 3 Å. Thus, the E484K, which is present in the Beta and Gamma variants of concern, and more recently in some Alpha isolates as well, is likely to have an enhancing role through transient salt bridges. The same cannot be said about E484Q seen in the Delta variant of concern because salt bridge formation is unlikely with the polar glutamine (Q) residue. With no accompanying aromatic change in positions 498 or 501, we believe that the E484Q in Delta, and additionally the K417N in ‘Delta Plus’, cannot sustain mouse infectivity on their own based on current biomolecular understanding. It also follows that the Omicron variant of concern is expected to infect wild type mice because it has the essential and enhancing mutations. The residues 75 to 82 in mACE2 are significantly different from hACE2 (**Table 3**), therefore any mutation in the corresponding RBD interface is worth investigation. We could find one from experimental observations, the engineered substitution F486L,^27^ and consider it to have at best an enhancing role. As it was observed simultaneously with Q498Y (which is likely to sustain mouse infectivity on its own), the contribution of F486L to the overall mouse adaptation remains to be ascertained.

### Comparison with *in vitro, in vivo* and *in situ* observations

Early *in silico* predictions based on comparative structural analysis of ACE2 suggested that mouse has a very low probability of being infected.^35,36^ Although correct about mouse, those analyses also made inconsistent and erroneous predictions that ferrets wouldn’t be susceptible, pigs would be susceptible, etc., thus exposing the need for experimental inputs into the model. Therefore, this paper takes into account a range of experimental observations to cross-check our *in silico* predictions through biomolecular modelling. Wan *et al*. reasoned that “*mouse or rat ACE2 contains a histidine at the 353 position which does not fit into the virus-receptor interaction as well as a lysine does*”. ^35^ While this is true of the original Wuhan strain containing asparagine 501 (N501) in the RBD, our modelling indicates why the tyrosine 501 mutation enables mouse infectivity, even on its own. In hACE2, lysine 353 (K353) creates a salt bridge with conserved aspartic acid 38 (D38). In mACE2, lysine 353 is replaced by the aromatic histidine (H353) to complete the salt bridge, as well as contribute to π-stacking with Y501 variant. The N501Y mutation will also lead to favorable π-stacking with the highly conserved tyrosine 41 (Y41) residue in mammalian ACE2, as suggested by Starr *et al*.^38^ with deep scanning of RBD mutations and hACE2 affinity assays. These authors highlighted enhanced affinity of F501 (as it had the highest score), followed by Y501, V501, W501 and T501, in that order. But Y501 and T501 require only a single nucleotide change and have been observed more frequently *in situ* (**Figure S1a**) – compared to F501, V501 and W501 which require 2, 2 and 3 nucleotide changes respectively. It is unsurprising that the latter variants requiring two or more changes were rarely observed *in situ* regardless of their high *in vitro* affinity scores from Starr *et al*.^38^

From the above and **Table 4**, we see that *in silico* analysis can provide valuable insights to interpret and bridge *in vitro, in vivo* and *in situ* observations on the RBD position 501. A similar analysis is possible with the alternative essential mutation for mouse adaptation at RBD position 498, where the *in vitro* affinity enhancement order is H498, Y498, F498 and W498 according to Starr *et al*.^38^ Of these, H498 (on its own) and Y498 (with enhancing RBD mutations) have been shown to result in mouse adaptation *in vivo* (**Table 1**);^7,8,23-28^ Q498R was also reported once, unusually in combination with N501Y, isolated from mouse lung after 30 passages. In humans, *in situ* observations of these variants have been limited to R498 (57 occurrences) and H498 (8 occurrences) so far. Thus, it is clear from *in vitro, in vivo* and *in situ* analyses (**Table 4**) that H498, R498 and Y498 are possible but not yet common. This is consistent with our *in silico* predictions because H498 and Y498 are aromatic (enabling π-stacking with ACE2 Y41; similar to Y501), while R498 has conformational flexibility and can still be associated with π-stacking interactions.^37^ H498 and R498 observed *in situ* require a single nucleotide change from Q498, while Y498 requires two nucleotide changes (or one change from H498).

**Table 4.**
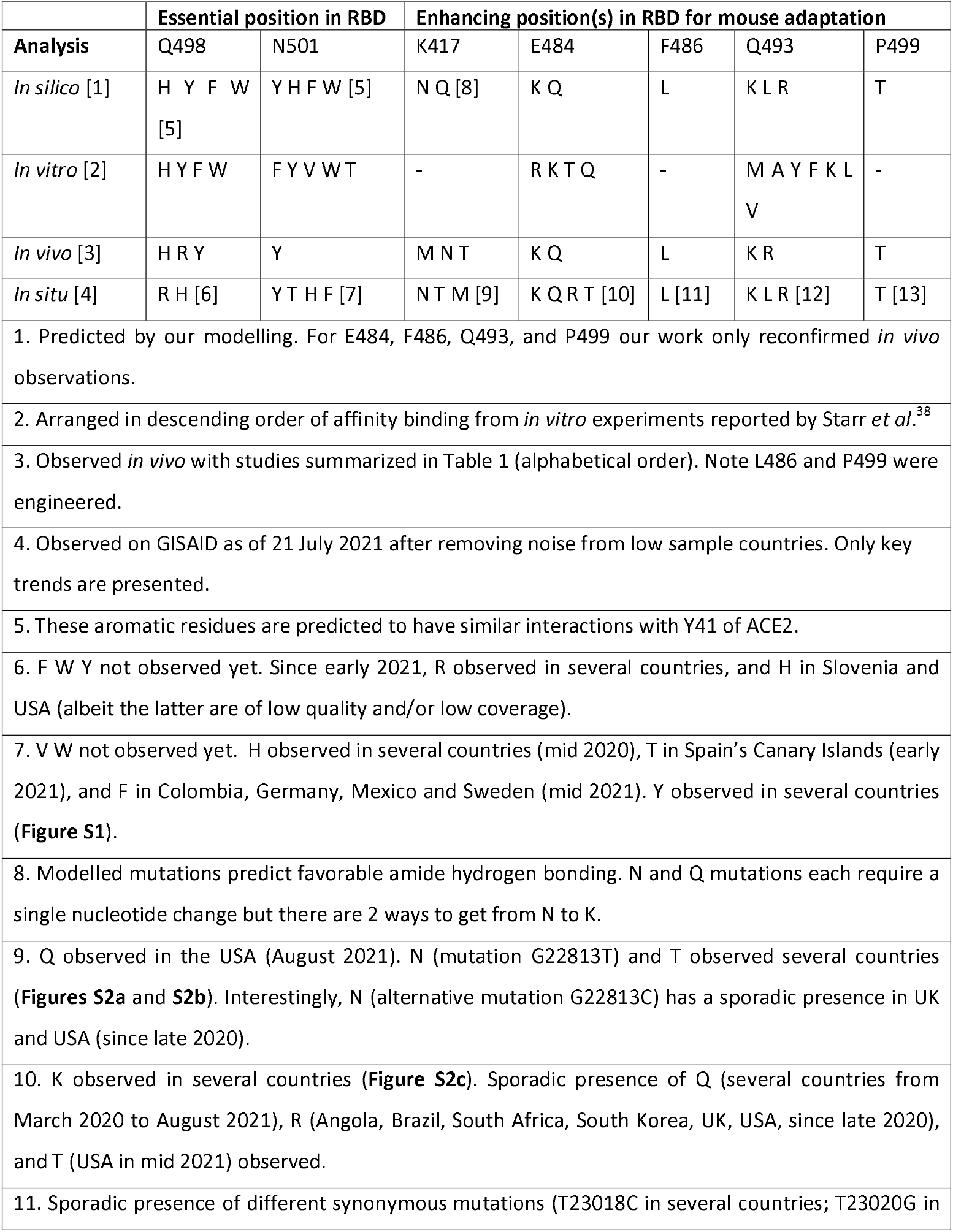
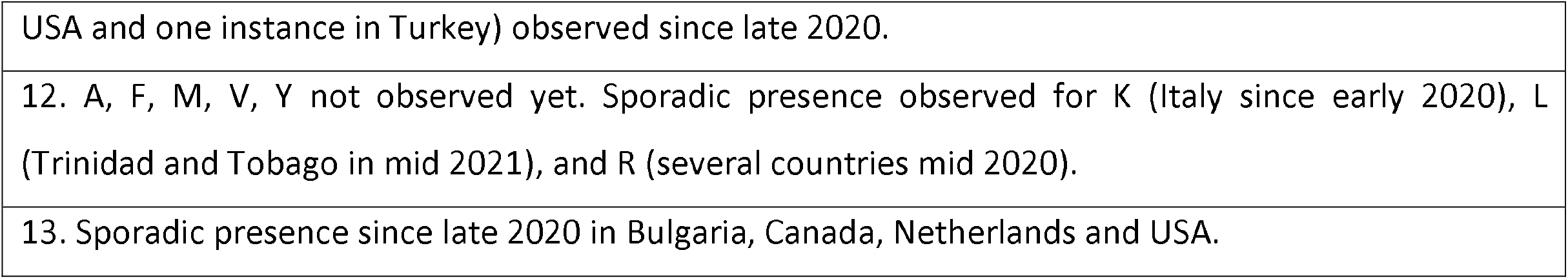
Comparison of *in silico, in vitro, in vivo* and *in situ* observations of key mutations in **Table 1**.

Similar insights are also possible for the enhancing RBD mutations (c.f. **Table 1** and **Table 4**). From *in vivo* and *in situ* observations, we see that K417M is less common than K417N or K417T (**Figures S2a** and **S2b**), although *in vitro* studies did not predict any enhancement.^38^ *In silico* predictions show that all these require a single nucleotide change, but that N417 (and Q417) would benefit from amide hydrogen bonding. E484K is present in Beta and Gamma (**Figure S2c**), and increasingly in Alpha VOC, while E484Q is present in the Kappa variant of interest that is related to the Delta VOC. In comparison to these two substitutions and notwithstanding higher *in vitro* affinity scores, R484 and T484 are infrequently observed *in situ*, which is consistent with our *in silico* predictions because they each require 2 nucleotide changes from E484, or one change from K484. The F486L and P499T were originally engineered *in vivo*, but have had sporadic *in situ* presence in human populations. *In silico* predictions suggest that the F486L mutation (accessible by three possible ways of a single nucleotide change) can aid mACE2 adaptation, due to the human-mouse differences in ACE2 at the 78-82 region; the P499T is also a single nucleotide change (but only one way from P to T) and predicted to be rare in comparison. Finally, *in silico* predictions for Q493 substituted by K, L or R (each a single nucleotide change) are borne out *in vivo* and *in situ*, although their affinity scores from *in vitro* experiments are low. The affinity scores from Starr *et al*.^38^ were developed for hACE2 (not mACE2), so we expected a greater correlation than what has been observed *in situ* in human populations, but perhaps it is still early in the pandemic to assess this definitively. It also looks likely that there are factors other than enhancement of ACE2 binding that determine susceptibility – as seen from Starr *et al*.^38^ and **Table 4**; from our own inconclusive attempts at correlating free energy binding affinities using *in silico* methods (not shown); and from a more detailed *in silico* model by Piplani *et al*.^*39*^ which counter-intuitively predicts lower binding free energy for dogs compared to more susceptible animals such as monkeys, hamsters, ferrets, cats and tigers.^40^

Some mutations in the essential as well as enhancing positions can lead to other mutations. For example, N501Y, the key mutation common to the Alpha, Beta, and Gamma variants of concern, can lead to F501 with a further nucleotide change. Similarly, the enhancing E484K mutation can also lead to R484 or T484 with a further nucleotide change. We examined whether *in situ* observations are consistent or contrary to our *in silico* predictions. Indeed, F501 was observed in Sweden (28 April 2021), Germany (7 May 2021), Mexico (22 June 2021) and Colombia (30 June 2021), once in each of these four countries, while the Y501 has been observed in these countries since 12 March 2020, 21 October 2020, 31 January 2021 and 19 September 2020 respectively. UK reported E484R in August 2020, followed by Angola in April 2021; Brazil and USA in May 2021; South Korea in June 2021; and South Africa in July 2021. In each case, the E484K was detected prior to E484R – the former mutation circulating in UK, Angola, Brazil, USA, South Korea and South Africa from April, August, April, March, December and August 2020 respectively. Similarly, E484T was only recently detected in the USA in June 2021, 15 months after the first report of E484K in that country. All eleven instances are thus consistent with our prediction – whether this link is causal or a coincidence is worthy of investigation with local epidemiological data. While bioinformatics tools can provide useful insights, out of 53 COVID-positive cases only one sample is on average sent for virus genome sequencing (as of 4 October 2021), with huge variations across time and locations, and lots of missing meta-data.^41,42^ This means we are more confident about ruling in (e.g. when a variant has been detected in a location) than ruling out the possibility of a mutation circulating purely based on *in silico* data, even if the latter is statistically large (4 million as of 4 October 2021).

Our analysis is not just of theoretical interest; it has huge practical applications because mice can be kept as pets, or come into contact with other pets like cats which are known to be susceptible. Also, mouse plague can occur in area of COVID-19 outbreaks or endemicity, as is currently the case in New South Wales and adjacent states of Australia.^43^ In order to help public health and animal health professionals, Figures 3a and 3b show the eleven countries where key essential and enhancing mutations listed in Tables 1 and 4 (viz. N501Y, E484K and K417N/T) have co-occurred. The underlying raw data, down to regional counts for these combinations, are available from https://www.biorxiv.org/content/10.1101/2021.08.04.455042v2.supplementary-material. We believe that this information will help locate areas at risk (especially in Brazil, Chile, Djibouti, Haiti, Malawi, Mozambique, Reunion, Suriname, Trinidad and Tobago, Uruguay and Venezuela) for appropriate mitigation measures.

**Figure 3.**
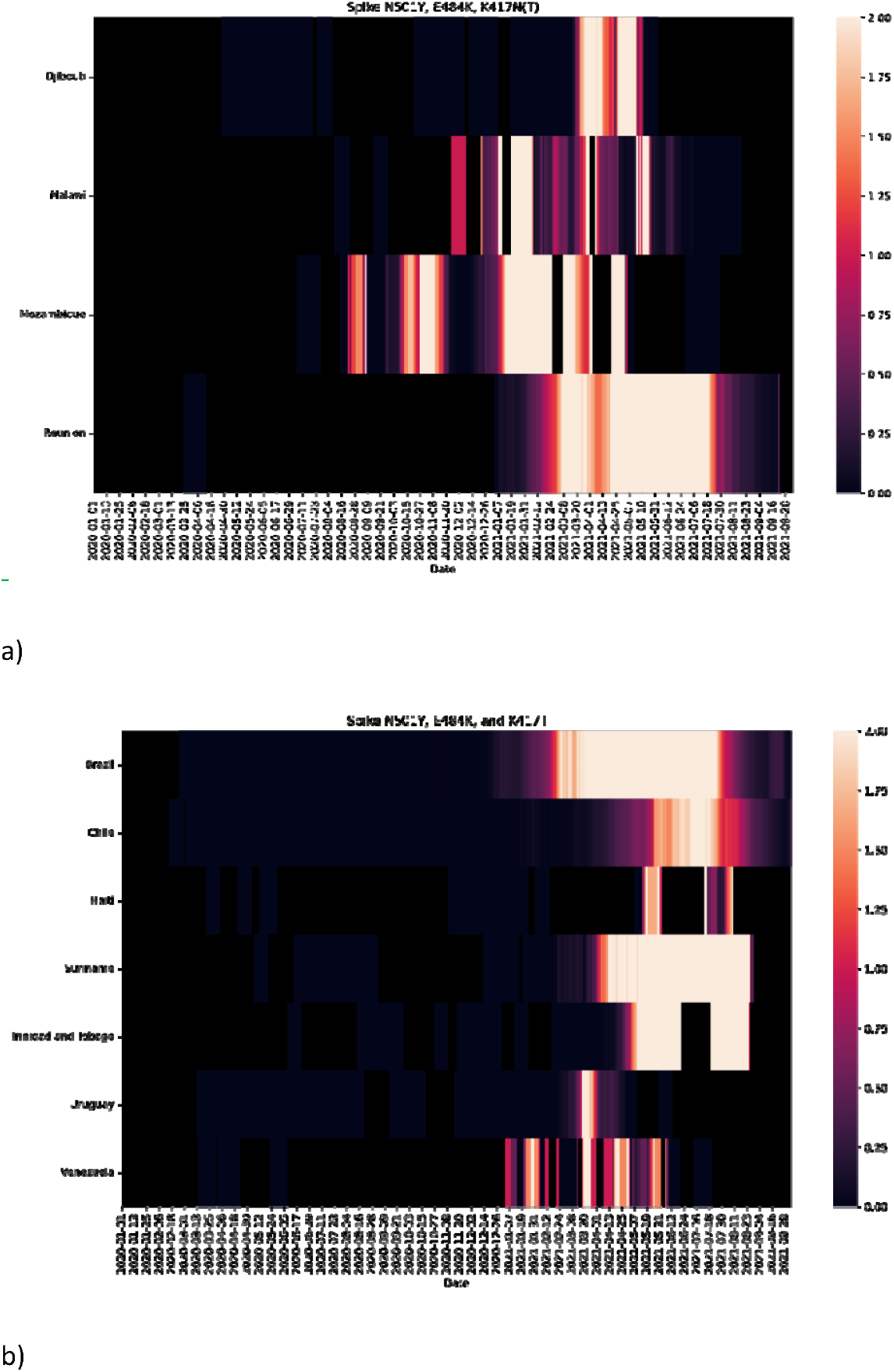
Significant occurrences on GISAID of **a)** N501Y, E484K and K417N (nucleotide change G22813T) triple mutations and **b)** N501Y, E484K and K417T triple mutations, at the virus Receptor Binding Domain (RBD) since the start of the COVID-19 pandemic (31 December 2019). These countries are encouraged to perform targeted field surveillance.

### Conclusion and further analyses

Assessing the risk of viruses adapting to new hosts requires careful interpretation of all available data from *in silico, in vivo, in vitro* and *in situ* sources. Understanding host adaptation at a molecular level, via modelling helps reconcile seemingly conflicting, experimental, and clinical observations while a pandemic is still in progress. We have demonstrated this with the SARS-CoV-2 virus adapting to mice. Our conclusions come with humility, as they are based on best available evidence up to this point, but allowing us and others to refine when more evidence becomes available. Armed with our collective understanding from different approaches, and bolstered by bioinformatics and emerging artificial intelligence technologies such as *AlphaFold*, we have shown how to position ourselves better to predict and mitigate virus host adaptations, not just for this pandemic but also for future Disease-X. Further analyses pertaining to COVID-19 should focus on the role of mutations beyond the spike and RBD; experimentally prove that the Omicron variant of concern can infect wild type mice; improve computational modelling of binding affinities to explore correlations with susceptibility, if any; assess the actual risk of transmission for this virus through aerosol versus other routes; and study other hosts like rats and other potential reservoir species (even those that previously exhibited low receptor activities) which will be hard to vaccinate or control.^35,36,44^

### Supplementary Methods for Molecular Modelling

Molecular simulations were performed using NAMD2.14^45^ with CHARM36m^46^ forcefield employing a ‘TIP3’ water model. The SARS-CoV-2 spike/ACE2 model was a homology model based on one of the best pdb structures available at the time of our analysis, *viz*. 6M17, which is deemed to be of sufficient quality for our purpose.^15^ Variant models of the SARS-CoV-2 spike RBD domain (residues 330 to 530) containing the mouse adapted mutations were constructed by mutating residues in the NAMD build scripts, but later found to be in very close agreement with the same model constructed using *AlphaFold*^*16*^ (less than 0.9 Å RMSD difference to C-Alpha backbone atoms). A truncated mACE2 consisting of residues 19 to 600 was built using Swiss modeller,^47^ and similarly found to be in close agreement with the equivalent *AlphaFold* model. Amino acid side chain conformations predicted by *AlphaFold* were used in all initial RBD model starting conformations. Glycosylation of the spike and mACE2 protein was manually constructed using Visual Molecular Dynamics (VMD). Simulations were run with Periodic Boundary Conditions ‘PBCs’ using the ‘NPT’ isothermal-isobaric ensemble at 310K and 1 bar pressure employing Langevin dynamics. The PBCs were constant in the XY dimensions. Long-range Coulomb forces were computed with the Particle Mesh Ewald method with a grid spacing of 1 Å. 2 fs timesteps were used with non-bonded interactions calculated every 2 fs and full electrostatics every 4 fs while hydrogens were constrained with the ‘SHAKE’ algorithm. The cut-off distance was 12 Å with a switching distance of 10 Å and a pair-list distance of 14 Å. Pressure was controlled to 1 atmosphere using the Nosé-Hoover Langevin piston method employing a piston period of 100 fs and a piston decay of 50 fs. Trajectory frames were captured every 100 ps. Eleven variant models were constructed representing the mouse-adapted variants observed in **Table 1** as well as the original Wuhan RBD model with both mouse and human ACE2. Models were simulated for 300 nanoseconds. Trajectories were visualized with VMD and Nanome. Modelling data shall be made available on the CSIRO Data access portal (https://data.csiro.au/).

## Acknowledgements

This work was supported by funding (Principal Investigator: S.S.V.) from the Australia’s Department of Finance, CSIRO Future Science Platforms, National Health and Medical Research Council (MRF2009092), and United States Food and Drug Administration (FDA) Medical Countermeasures Initiative contract (75F40121C00144). The article reflects the views of the authors and does not represent the views or policies of the funding agencies including the FDA. We are grateful for support from our colleagues at the Australian Centre for Disease Preparedness (https://www.grid.ac/institutes/grid.413322.5) (especially Simran Chahal, Trevor Drew, Alexander McAuley and Nagendrakumar Singanallur) and the Transformational Bioinformatics Group (especially Denis Bauer, Yatish Jain, Brendan Hosking and Aidan Tay). L.O.W.W acknowledges grant funding from the Australian Academy of Science and Australia’s Department of Industry, Science, Energy and Resources. The title is from the poem ‘*To a Mouse: On Turning her up in her Nest, with the Plough, November 1785*’ by Scotland’s national poet Robert Burns, in which he says that the mouse is not alone in proving foresight may be vain as the best-laid schemes of mice and men go oft awry (*But, Mousie, thou art no thy-lane, In proving foresight may be vain: The best-laid schemes o’ Mice an’ Men Gang aft agley*).

## Author contributions

Conceptualization, methodology, and funding acquisition, S.S.V.; *in silico* analysis, M.J.K.; *in vitro* analysis, M.J.K. and S.S.V.; *in vivo* analysis, S.M. and S.S.V.; *in situ* analysis, L.O.W.W., D.R. and C.L.; writing – original draft preparation, S.S.V, M.J.K. and S.M.; writing – review and editing, all authors.

---

**Figure S1.**
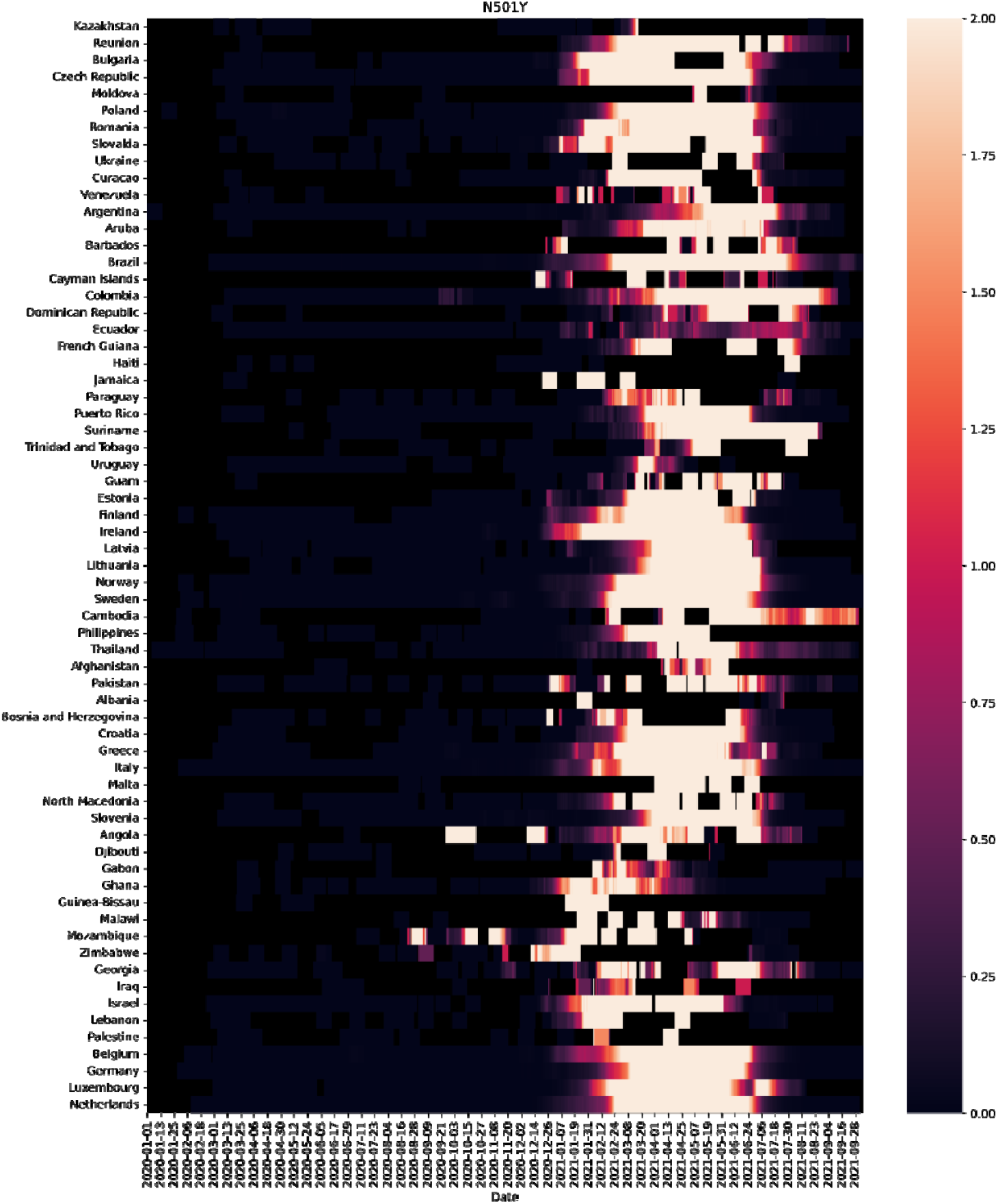
Significant occurrences on GISAID of the essential N501Y mutation (nucleotide change A23063T), at the virus Receptor Binding Domain (RBD) since the start of the COVID-19 pandemic (31 December 2019).

**Figure S2.**
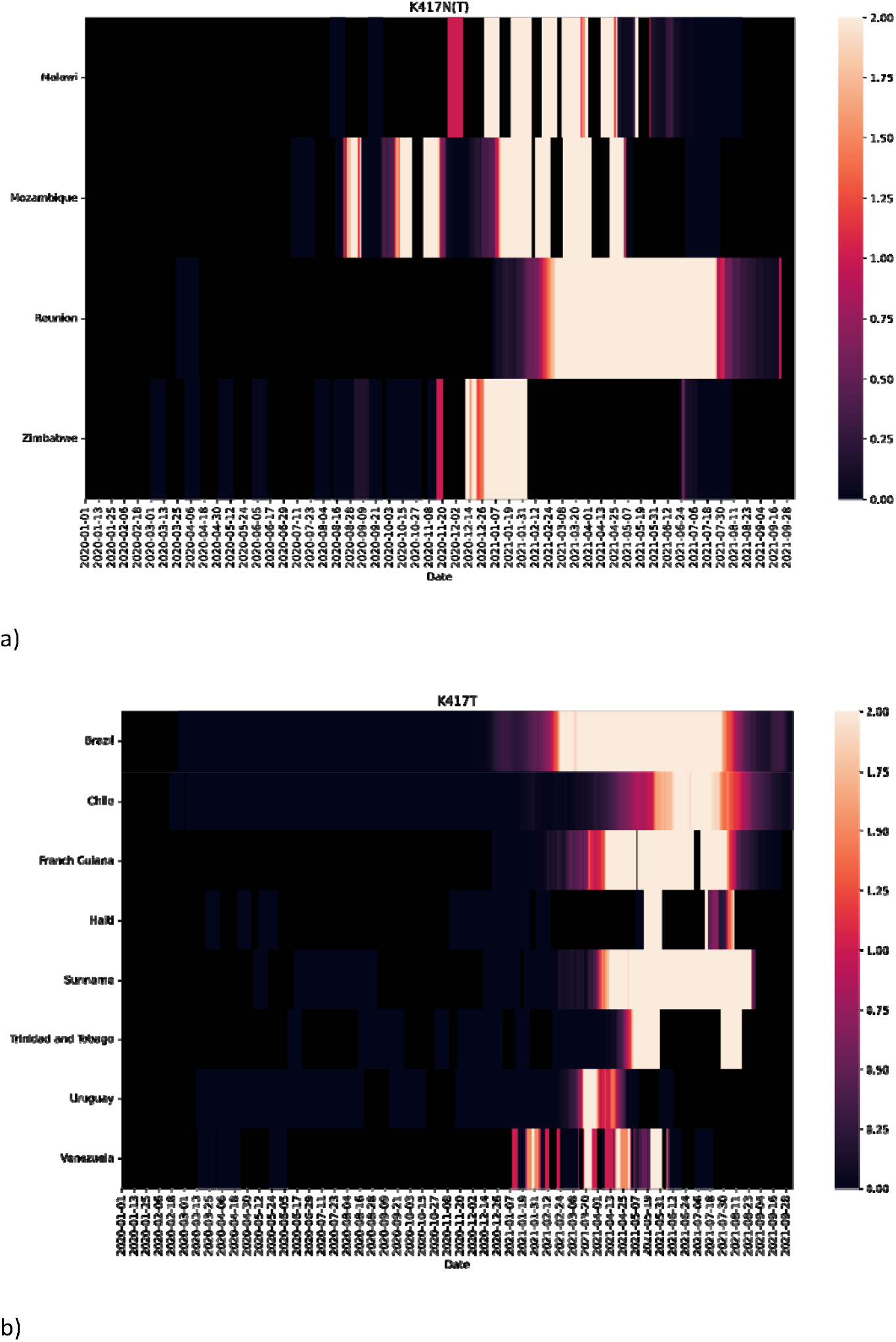
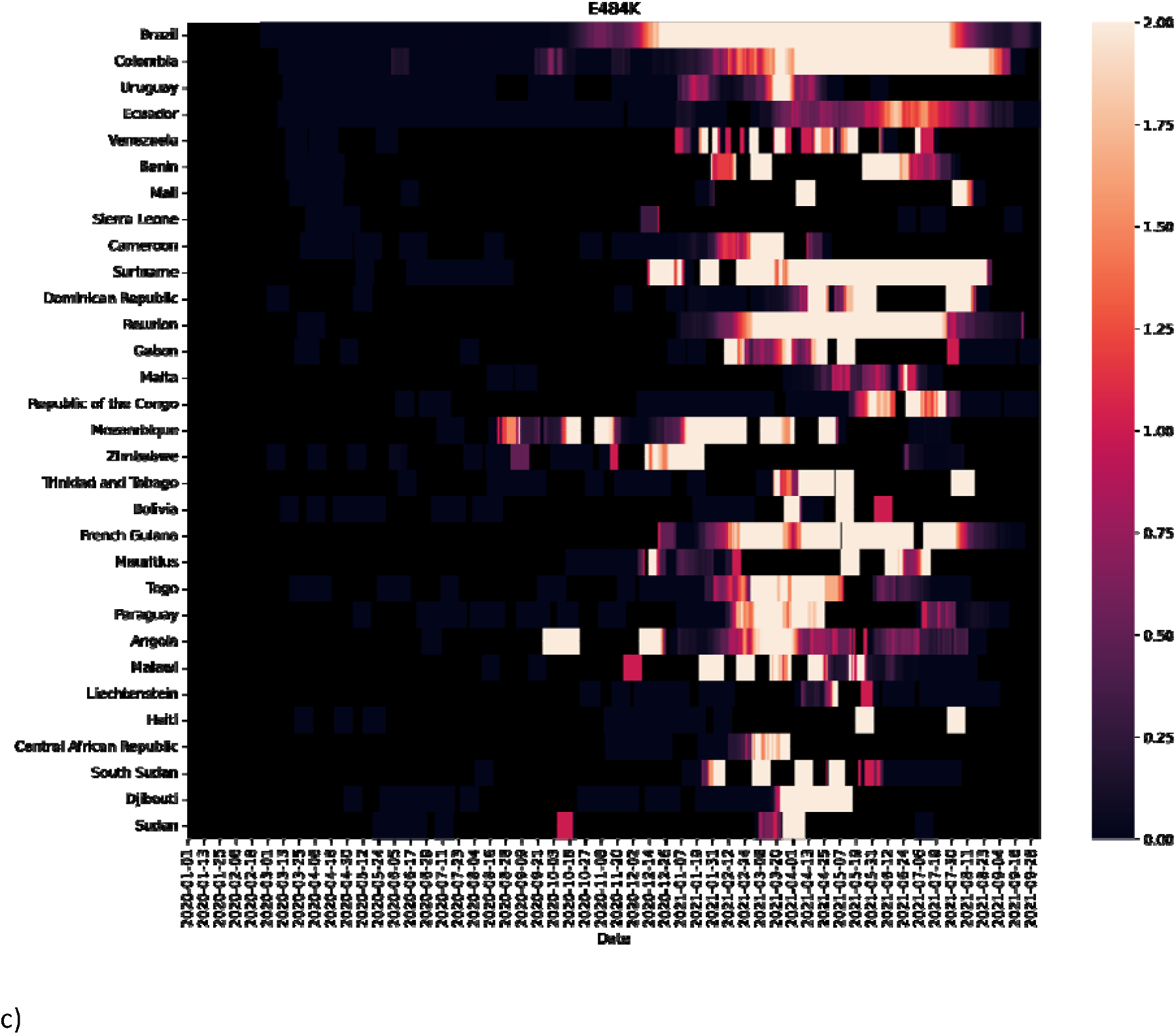
Significant occurrences on GISAID of the enhancing mutations **a)** K417N (nucleotide change G22813T), **b)** K417T (nucleotide change A22812C), **c)** E484K (nucleotide change G23012A), at the virus Receptor Binding Domain (RBD), since the start of the COVID-19 pandemic (31 December 2019).

## References

1. Dong E, Du H, Gardern L. An interactive web-based dashboard to track COVID-19 in real time. Lancet Infectious Diseases. 2020;20(5):533–534. Available at https://doi.org/10.1016/S1473-3099(20)30120-1 (accessed 1 December 2020).

2. Bauer DC, Tay AP, Wilson, LOW, et al. Supporting pandemic response using genomics and bioinformatics: A case study on the emergent SARS-CoV-2 outbreak. Transboundary & Emerging Diseases. 2020; 67(4):1453–1462. Available at https://doi.org/10.1111/tbed.13588 (accessed 1 December 2020).

3. Lassaunière R, Fonager J, Rasmussen, M, et al. Working paper on SARS-CoV-2 spike mutations arising in Danish mink, their spread to humans and neutralization data. 2020. Available at https://files.ssi.dk/Mink-cluster-5-short-report_AFO2 (accessed 1 December 2020).

4. Korber B, Fischer WM, Gnanakaran S, et al. Tracking changes in SARS-CoV-2 spike: Evidence that D614G increases infectivity of the COVID-19 virus. Cell. 2020; 182(4):812–827. Available at https://doi.org/10.1016/j.cell.2020.06.043 (accessed 1 December 2020).

5. Zhang J, Cai Y, Xiao T, et al. Structural impact on SARS-CoV-2 spike protein by D614G substitution. Science. 2021; 372(6541):525–530. Available at https://doi.org/10.1126/science.abf2303 (accessed 1 May 2021).

6. Gu H, Chen Q, Yang G, et al. Adaptation of SARS-CoV-2 in BALB/c mice for testing vaccine efficacy. Science. 2020; 369(6511):1603–1607. Available at https://doi.org/10.1126/science.abc4730 and https://www.biorxiv.org/content/10.1101/2020.05.02.073411v1 (accessed 1 December 2020).

7. Dinnon KH, Leist SR, Schäfer A, et al. A mouse-adapted model of SARS-CoV-2 to test COVID-19 countermeasures. Nature. 2020; 586:560–566. Available at https://doi.org/10.1038/s41586-020-2708-8 (accessed 1 December 2020).

8. Leist SR, Dinnon KH, Schäfer A, et al. A mouse-adapted SARS-CoV-2 induces acute lung injury and mortality in standard laboratory mice. Cell. 2020; 183:1070–1085. Available at https://doi.org/10.1016/j.cell.2020.09.050 (accessed 1 December 2020).

9. Yuan L, Tang Q, Cheng T, Xia N. Animal models for emerging coronavirus: progress and new insights. Emerging Microbes & Infections. 2020; 9(1):949–961. Available at https://doi.org/10.1080/22221751.2020.1764871 (accessed 15 May 2021).

10. Muñoz-Fontela C, Dowling WE, Funnell SGP, et al. Animal models for COVID-19. Nature. 2020; 586: 509–515. Available at https://doi.org/10.1038/s41586-020-2787-6 (accessed 1 December 2020).

11. Zeiss CJ, Compton S, Veenhuis, RT. Animal models of COVID-19. I. Comparative virology and disease pathogenesis. ILAR Journal. 2021; ilab007. Available at https://doi.org/10.1093/ilar/ilab007 (accessed 1 May 2021).

12. Veenhuis RT, Zeiss CJ. Animal models of COVID-19 II. Comparative immunology. ILAR Journal. 2021; ilab010. Available at https://doi.org/10.1093/ilar/ilab010 (accessed 1 May 2021).

13. Rathnasinghe R, Jangra S, Cupic A, et al. The N501Y mutation in SARS-CoV-2 spike leads to morbidity in obese and aged mice and is neutralized by convalescent and post-vaccination human sera. medRxiv. 2021; 2021.01.19.21249592. Available at https://doi.org/10.1101/2021.01.19.21249592 (accessed 1 February 2021).

14. Yao W, Wang Y, Ma D, et al. Circulating SARS-CoV-2 variants B.1.1.7, 501Y.V2, and P.1 have gained ability to utilize rat and mouse Ace2 and altered in vitro sensitivity to neutralizing antibodies and ACE2-Ig. bioRxiv. 2021; 2021.01.27.428353. Available at https://doi.org/10.1101/2021.01.27.428353 (accessed 1 May 2021).

15. Yan R, Zhang Y, Li Y, et al. Structural basis for the recognition of SARS-CoV-2 by full-length human ACE2. Science. 2020; 367:1444–1448. Available at https://doi.org/10.1126/science.abb2762 (accessed 1 December 2020).

16. Jumper, J., Evans, R., Pritzel, A. et al. Highly accurate protein structure prediction with AlphaFold. Nature (2021). Available at https://doi.org/10.1038/s41586-021-03819-2 (accessed 20 July 2021).

17. McAuley AJ, Kuiper MJ, Durr PA, et al. Experimental and in silico evidence suggests vaccines are unlikely to be affected by D614G mutation in SARS-CoV-2 spike protein. Npj Vaccines. 2020; 5:96. Available at https://doi.org/10.1038/s41541-020-00246-8 (accessed 1 December 2020).

18. Riddell S, Goldie S, McAuley A, et al. Live virus neutralisation of the 501Y.V1 and 501Y.V2 SARS-CoV-2 variants following INO-4800 vaccination of ferrets. bioRxiv. 2021; 2021.04.17.440246. Available at https://doi.org/10.1101/2021.04.17.440246 (accessed 1 May 2021).

19. Tai W, He L, Zhang X. Characterization of the receptor-binding domain (RBD) of 2019 novel coronavirus: implication for development of RBD protein as a viral attachment inhibitor and vaccine. Cellular & Molecular Immunology. 2020; 17:613–620. Available at https://doi.org/10.1038/s41423-020-0400-4 (accessed 1 December 2020).

20. Grantham, R. Amino acid difference formula to help explain protein evolution. Science. 1974; 185(4154): 862–864. Available at https://doi.org/10.1126/science.185.4154.862 (accessed 1 December 2020).

21. Sun S, Gu H, Cao L, et al. Characterization and structural basis of a lethal mouse-adapted SARS-CoV-2. Nature Communications. 2021; 12: 5654. Available at https://www.nature.com/articles/s41467-021-25903-x (accessed 30 September 2021).

22. Fagre A, Lewis J, Eckley M, et al. SARS-CoV-2 infection, neuropathogenesis and transmission among deer mice: Implications for spillback to New World rodents. PLoS Pathogens. 2021; 17(5): e1009585. Available at https://doi.org/10.1371/journal.ppat.1009585 (accessed 24 May 2021).

23. Wang J, Shuai L, Wang C, et al. Mouse-adapted SARS-CoV-2 replicates efficiently in the upper and lower respiratory tract of BALB/c and C57BL/6J mice. Protein & Cell. 2020; 11:776–782. Available at https://doi.org/10.1007/s13238-020-00767-x (accessed 1 December 2020).

24. Liu Z, Zheng H, Lin H, et al. Identification of common deletions in the spike protein of Severe Acute Respiratory Syndrome Coronavirus 2. Journal of Virology. 2020; 94(17):e00790–20. Available at https://doi.org/10.1128/JVI.00790-20 (accessed 1 December 2020).

25. Zhou H, Chen X, Hu T, et al. A novel bat coronavirus closely related to SARS-CoV-2 contains natural insertions at the S1/S2 cleavage site of the spike protein. Current Biology. 2020; 30:2196–2203. Available at https://doi.org/10.1016/j.cub.2020.05.023 (accessed 1 December 2020).

26. Huang K, Zhang Y, Hui X, et al. Q493K and Q498H substitutions in spike promote adaptation of SARS-CoV-2 in mice. EbioMedicine. 2021; 67:103381. Available at https://doi.org/10.1016/j.ebiom.2021.103381 (accessed 24 May 2021).

27. Muruato A, Vu MN, Johnson BA, et al. Mouse adapted SARS-CoV-2 protects animals from lethal SARS-CoV challenge. bioRxiv. 2021; 2021.05.03.442357. Available at https://doi.org/10.1101/2021.05.03.442357 (accessed 15 May 2021).

28. Zhang Y, Huang K, Wang T, et al. SARS-CoV-2 rapidly adapts in aged BALB/c mice and induces typical pneumonia. Journal of Virology. 2021; 95(11):e02477–20. Available at https://doi.org/10.1128/JVI.02477-20 (accessed 24 May 2021).

29. Van Noorden, R. Scientists call for fully open sharing of coronavirus genome data. Nature. 2021; 590:195–196. Available at https://doi.org/10.1038/d41586-021-00305-7 (accessed 1 March 2021).

30. Rambaut A, Loman N, Pybus O, et al. Preliminary genomic characterisation of an emergent SARS-CoV-2 lineage in the UK defined by a novel set of spike mutations. Virological. 2020. Available at https://virological.org/t/preliminary-genomic-characterisation-of-an-emergent-sars-cov-2-lineage-in-the-uk-defined-by-a-novel-set-of-spike-mutations/563 (accessed 1 March 2021).

31. Li Q, Nie J, Wu J, et al. SARS-CoV-2 501Y.V2 variants lack higher infectivity but do have immune escape. Cell. 2021; 184(9):2362–2371. Available at https://doi.org/10.1016/j.cell.2021.02.042 (accessed 1 May 2021).

32. Montagutelli X, Prot M, Levillayer L, et al. The B1.351 and P.1 variants extend SARS-CoV-2 host range to mice. bioRxiv. 2021; 2021.03.18.436013. Available at https://doi.org/10.1101/2021.03.18.436013 (accessed 1 May 2021).

33. Faria NR, Claro IM, Candido D, et al. Genomic characterisation of an emergent SARS-CoV-2 lineage in Manaus: preliminary findings. Virological. 2021. Available at https://virological.org/t/genomic-characterisation-of-an-emergent-sars-cov-2-lineage-in-manaus-preliminary-findings/586 (accessed 1 March 2021).

34. Lucaci AG, Zehr JD, Shank SD, et al. RASCL: Rapid assessment of SARS-COV-2 clades enabled through molecular sequence analysis and its application to B.1.617.1 and B.1.617.2. Virological. 2021. Available at https://virological.org/t/rascl-rapid-assessment-of-sars-cov-2-clades-enabled-through-molecular-sequence-analysis-and-its-application-to-b-1-617-1-and-b-1-617-2/709 (accessed 29 May 2021).

35. Wan Y, Shang J, Graham R, et al. Receptor recognition by novel coronavirus from Wuhan: an analysis based on decade-long structural studies of SARS coronavirus. Journal of Virology. 2020; 94(7):e00127–20. Available at https://doi.org/10.1128/JVI.00127-20 (accessed on 1 December 2020).

36. Damas J, Hughes GM, Keough C, et al. Broad host range of SARS-CoV-2 predicted by comparative and structural analysis of ACE2 in vertebrates. PNAS. 2020; 117(36):22311–22322. Available at https://doi.org/10.1073/pnas.2010146117 (accessed 1 December 2020).

37. Flocco MM, Mowbray SL. Planar stacking interactions of arginine and aromatic side-chains in proteins. Journal of Molecular Biology. 1994; 235(2):709–717. Available at https://doi.org/10.1006/jmbi.1994.1022 (accessed 1 December 2020).

38. Starr TN, Greaney AJ, Hilton SK, et al. Deep mutational scanning of SARS-CoV-2 receptor binding domain reveals constraints on folding and ACE2 binding. Cell. 2020; 182(5):1295–1310.e20. Available at https://doi.org/10.1016/j.cell.2020.08.012 (accessed 1 December 2020).

39. Piplani S, Singh PK, Winkler DA, Petrovsky, N. In silico comparison of SARS-CoV-2 spike protein-ACE2 binding affinities across species and implications for virus origin. Scientific Reports. 2021; 11:13063. Available at https://doi.org/10.1038/s41598-021-92388-5 (accessed 1 October 2021).

40. Muñoz-Fontela C, Dowling, WE, Funnell, SGP, et al. Animal models for COVID-19. Nature. 2020; 586:509–515. Available at https://doi.org/10.1038/s41586-020-2787-6 (accessed 1 December 2020).

41. Bauer DC, Metke-Jimenez A, Maurer-Stroh S, et al. Interoperable medical data: the missing link for understanding COVID-19. Transboundary & Emerging Diseases. 2021; 68(4):1753–1760. Available at https://onlinelibrary.wiley.com/doi/10.1111/tbed.13892 (accessed 1 July 2021).

42. Priyadarshini, S. Massive coronavirus sequencing efforts urgently need patient data. Nature India (special issue #13 on COVID-19 crisis). 2020; 10.1038/nindia.2020.75:11-13. Available at https://www.natureasia.com/en/nindia/pdf/special-issues/13/Nature-India-COVID-19-Crisis.pdf and https://go.nature.com/2y7kUIw (accessed 1 December 2020).

43. NSW Government. Help for regional communities impacted by the mouse plague.2021. Available at https://www.nsw.gov.au/initiative/mouse-control-support-program (accessed 24 May 2021).

44. Zhao X, Chen D, Szabla R, et al. Broad and differential animal angiotensin-converting enzyme 2 receptor usage by SARS-CoV-2. Journal of Virology. 2020; 94(18):e00940–20. Available at https://doi.org/10.1128/JVI.00940-20 (accessed 1 December 2020).

45. Phillips JC, Braun R, Wang W, et al. Scalable molecular dynamics with NAMD. Journal of Computational Chemistry. 2005; 26:1781–1802. Available at https://doi.org/10.1002/jcc.20289 (accessed 1 December 2020).

46. Huang J, Rauscher S, Nawrocki G, et al. CHARMM36m: an improved force field for folded and intrinsically disordered proteins. Nature Methods. 2017; 14:71–73. Available at: https://doi:10.1038/nmeth.4067 (accessed 1 December 2020).

47. Waterhouse A, Bertoni M, Bienert S, et al. SWISS-MODEL: homology modelling of protein structures and complexes. Nucleic Acids Research. 2018; 46:W296–W303. Available at https://doi.org/10.1093/nar/gky427 (accessed 1 December 2020).

